# Cardiac-phase gating of eye movements: evidence from a gaze-contingent “searchlight” paradigm

**DOI:** 10.1101/2025.05.23.655692

**Authors:** Akane Hisada, Tomoko Isomura

## Abstract

Humans actively sample their environment by alternating rapid saccades for exploration with fixations for detailed perception, as high-resolution vision is restricted to the fovea. Efficient visual sampling depends on the precise coordination of these eye movements. Here, we show that this coordination is gated by the cardiac cycle. Using a gaze-contingent “searchlight” paradigm, we limited visibility to a foveal window—thereby inducing frequent saccades and clearly dissociating the roles of saccades and fixations during natural-scene exploration—while simultaneously recording ECG. We found that saccades occur predominantly during systole, when baroreceptors are active, whereas fixations predominate during diastole, when baroreceptor-mediated afferent signals are quiescent. These findings identify the heartbeat as an internal clock that choreographs exploration and perception, revealing a mechanism through which interoceptive signals organize spontaneous perception–action cycles.

## 1. Introduction

Human perception is inherently dynamic, characterized by continuous fluctuations arising from both spontaneous neural activity and the regular rhythmic patterns of our visceral organs. These inherent variations are not merely background noise, but are fundamental to sensory information processing and integration. In particular, the rhythmic activity of the visceral organs—interoceptive rhythms—has garnered increasing attention as it appears to play a critical role in shaping our perceptual experiences (Engelen et al., 2023). Among the various systems that generate interoceptive rhythms— such as those of the heart, respiration, and gastric activity—cardiac activity has received the most attention.

Research on the human cardiac cycle has shown that perception and cognition are modulated by its phases. This modulation has been particularly studied by presenting exteroceptive stimuli—such as visual, auditory, tactile, and pain—in synchrony with distinct cardiac phases (for review, see Garfinkel & Critchley, 2016; Skora et al., 2022). The method of presenting stimuli synchronized with specific phases of the cardiac cycle has been groundbreaking and has significantly advanced interoceptive research; however, it is important to note that in everyday life we are not passively exposed to stimuli in this manner. Rather, we actively engage in perceiving the world by moving and adjusting our sensory organs. For visual perception, we deliberately move our eyes to direct objects onto the fovea—the region of the retina where visual receptors are packed most densely—to optimize image resolution. This suggests that rhythmic oscillations of the cardiac cycle might modulate the way we actively sample visual information from the world. A recent study showed that saccades and fixations are significantly coupled with the cardiac cycle during a “spot the difference” visual search task (Galvez-Pol et al., 2020). Building on these findings, we further advanced the approach to enhance the ecological validity of the task and highlighted its unique contribution to the understanding of how the cardiac cycle influences active visual sampling.

In this study, we developed an original “searchlight task” that required participants to actively sample visual information by deliberately moving their eyes. In this task, participants viewed natural scene images, with only the central visual field visible while peripheral regions were masked in black using real-time gaze-contingent manipulation. By blocking peripheral input, we created a scenario in which participants must execute eye movements to gather the necessary information, thereby clearly differentiating the functional roles of fixations and saccades. The task was self-paced, allowing participants to engage in motivation-driven visual scanning. ECG signals were recorded during the task to test the coupling between cardiac phase and eye movements. We hypothesized that fixations—periods during which visual information is sampled—would occur more frequently during diastole, whereas saccades—which transiently impair visual perception but are essential for directing gaze toward the fovea—would occur more frequently during systole.

## 2. Results

A total of 41 participants performed the searchlight task, reporting the content of each image. Because only the central visual field was visible, participants had to actively move their eyes to scan the entire image, much like a searchlight (see Methods). Each participant completed 21 trials, each lasting 40 seconds. ECG was recorded simultaneously, and we computed the coupling between cardiac phase and ocular behaviors, including saccades, fixations, and blinks (Fig. 1B). The mean R-R interval across participants during the task was 819 ms (SD = 32.12, range: 691–942). During each 40-second trial, participants made an average of 28 saccades (SD = 5.19, range: 17–46), 44 fixations (SD = 12.22, range: 21–69), and 8 blinks (SD = 4.80, range: 2–19).

**Fig. 1.**
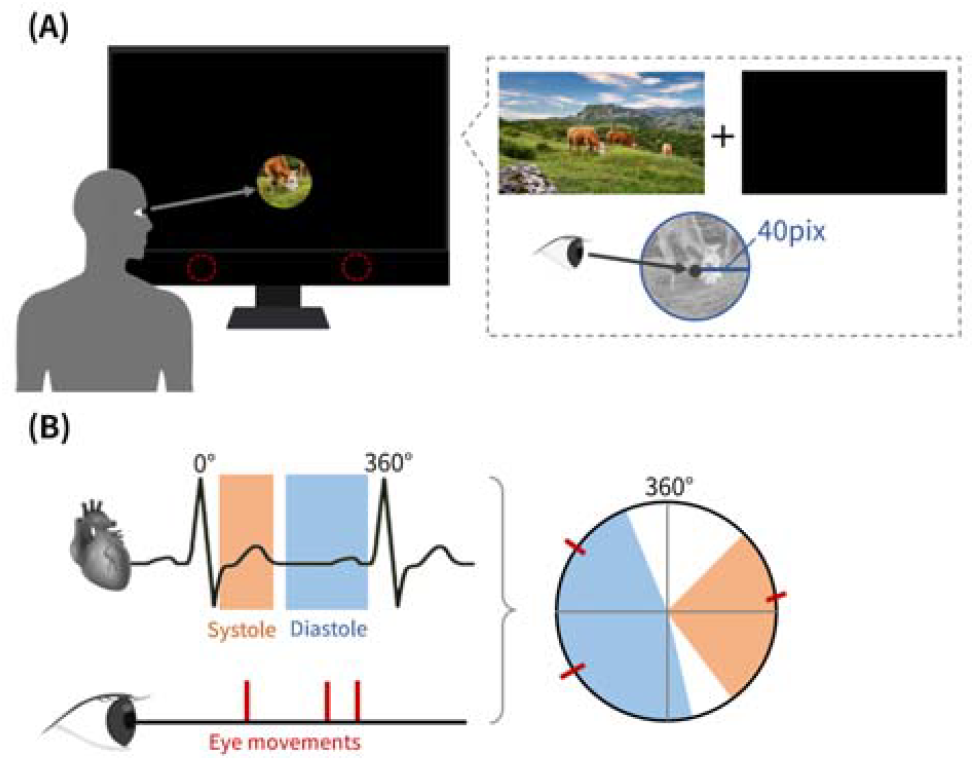
Schematic diagram of the task and analysis. (A) Example of a single trial. Participants were required to report the content of stimulus images hidden behind black masks. As they moved their eyes to scan the stimulus images on the display, a portion of the stimulus image became visible within a circular area (40-pixel radius) centered on their gaze position. (B) Schematic overview of the analysis. The top-left panel shows the ECG waveform, with the systole phase indicated in orange and the diastole phase in blue. The bottom-left panel shows the timing of eye movements, indicated by red lines. The right panel shows a clockwise circle with R-peaks set to 0 degrees. The timing of eye movements is plotted as red lines along the circumference.

To test the hypothesis that eye movements are coupled with cardiac cycles, we employed circular statistics, which account for the cyclical nature of the cardiac rhythm. For each participant, the median phase (in degrees) of saccades, fixations, and blinks within the R-R interval was calculated and used as the representative value.

At the group level, saccades occurred on average at 119.83° within the R-R interval, as indicated by the red arrow in Fig. 2A. To test whether the distribution was skewed toward a specific cardiac phase, we performed a Rayleigh test with the null hypothesis of uniformity. The results showed a significant deviation from uniformity (z = 0.34, p = 0.009), with the peak at 119.83° corresponding to the early stage of the cardiac cycle—the systolic phase (Fig. 2A).The same analysis was conducted for fixations and blinks. Fixations occurred on average at 204.32° within the R-R interval, significantly deviating from uniformity (z = 0.33, p = 0.013), corresponding to the mid-phase—the early diastolic phase (Fig. 2B). In contrast, blinks occurred on average at 297.31°. Although the Rayleigh test for blinks was not statistically significant (z = 0.26, p = 0.063), it showed a trend toward non-uniformity, suggesting that blink timing corresponded to the late period of the cardiac cycle—the diastolic phase (Fig. 2C).

**Fig. 2.**
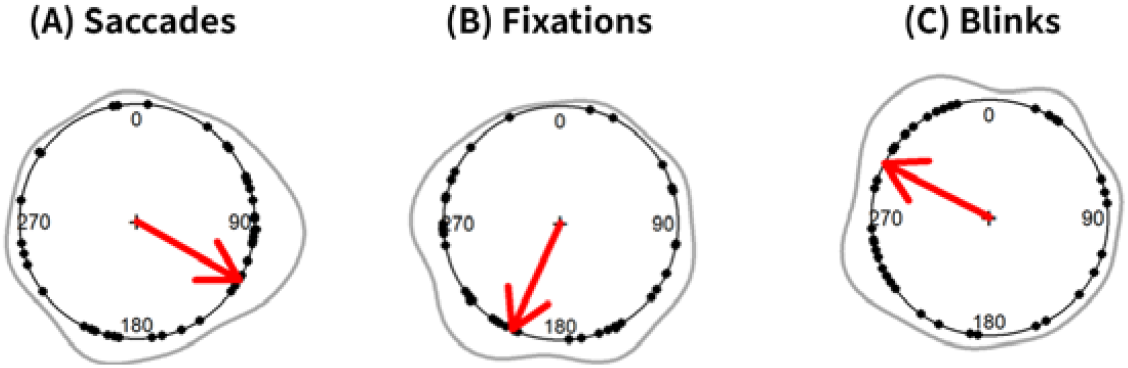
Relationship between eye movement timing and the cardiac cycle. In each panel, the timing of eye movements is plotted on a circular scale with the R-peaks as the vertex. Black dots indicate the median timing of saccades, fixations, and blinks for each participant across all trials. Red lines indicate phase bias vectors. Gray wavy lines represent the kernel density estimation. (A), (B), and (C) indicates the relationships between the timing of saccades, fixations, and blinks, respectively, and the phase of the R-R interval.

## 3. Discussion

Humans actively explore their environment by moving their sensory organs to gather information. In vision, we incessantly shift our gaze because only the fovea—a small retinal region densely packed with cone photoreceptors—supports high-acuity color vision. To place an object of interest onto the fovea, the oculomotor system generates rapid ballistic eye movements known as saccades. Although such movements should blur incoming input, perceptual stability is preserved by saccadic suppression, a predictive inhibitory mechanism by which the brain transiently attenuates visual processing around each saccade. Perception resumes as soon as the eye comes to rest in the ensuing fixation (Hamker et al., 2008; Ibbotson et al., 2011; Poletti et al., 2013).

The efficient operation of this system benefits from the coordinated timing of saccades and fixations. In this study, we show that the cardiac cycle gates active visual sampling. Eye movement timing was coupled with the cardiac cycle: saccades occurred predominantly during systole, whereas fixations predominated during diastole. These findings suggest that the cardiac cycle acts as an internal clock that choreographs when we explore and when we perceive.

Each heartbeat triggers baroreceptors in the aortic arch and carotid sinus, sending a burst of afferent activity to the brainstem and insular networks that maintain cardio-visceral homeostasis (Critchley & Harrison, 2013; Khalsa et al., 2018; Engelen et al., 2023). Similar to the blur induced by saccades, this baroreceptor volley can act as an internal noise that competes with the incoming exteroceptive information. For stable perception of the world, the neural predictive processes would need to suppress sensory gain during systole, the baroreceptor-active phase, and release that suppression during diastole.

Within a predictive-coding framework, the heartbeat entrains cortical rhythms such that precision—the confidence assigned to prediction errors—is rhythmically weighted. During systole, high interoceptive precision biases processing toward self-related bodily signals and away from external inputs, creating a favorable window for action initiation, which also generates self-originating somatosensory feedback that should be damped. During diastole, interoceptive precision drops, exteroceptive channels are up-weighted, and perceptual sampling is enhanced (Skora et al., 2022). In this view, the cardiac cycle dynamically toggles the brain between “action-centered” and “perception-centered” modes, explaining why we observed saccade clustering in systole and fixation-based information gathering in diastole.

Converging behavioral evidence supports this heartbeat-gated principle across diverse active tasks, including saccades during a spot-the-difference visual-search paradigm (Galvez-Pol et al., 2020); microsaccades during stable fixations (Ohl et al., 2016); self-paced button presses in a memory-recall task (Kunzendorf et al., 2019). Our findings extend the literature to a more ecologically valid form of visual exploration and further strengthen the case a heartbeat-locked mechanisms that coordinates action with perception.

In sum, this study is the first to show cardiac modulation of active visual sampling during natural-scene viewing, using a novel gaze-contingent “searchlight” paradigm that exaggerates the fovea–periphery contrast. This approach opens a path to studying how interoception orchestrates spontaneous perception–action cycles under ecologically valid conditions. Because the task is non-verbal, it can be applied to infants, clinical populations, or animals, offering a powerful tool for probing the biological basis—and potentially the causal architecture—of interoceptive–exteroceptive coupling across the lifespan.

## 4. Materials and methods

### 4.1 Participants

The final sample consisted of 41 healthy adults (25 females; mean age = 21.0 years, SD = 1.93), excluding one due to an error in the ECG recording. Participants had normal vision (corrected or uncorrected) and no known cardiac abnormalities. This study was approved by the Ethics Committee of the Department of Psychology at Nagoya University (approval number: NUPSY -240305-L-01), and informed consent was obtained from all participants.

### 4.2 Stimulus and Apparatus

The stimulus images (26 in total) were selected from Pixabay, a royalty-free repository. These images contained information about animals and their backgrounds (e.g., three goats in a meadow) and were edited to a resolution of 1920 (W) × 1282 (H) pixels. The task was presented on a 24.5-inch LCD monitor (240 Hz refresh rate) and controlled by a custom-written script in MATLAB R2019b (MathWorks, Inc.) with Psychophysics Toolbox extensions (Brainard, 1997; Pelli, 1997; Kleiner et al., 2007). A Tobii TX300 eye tracker (Tobii AB) was positioned below the monitor, allowing us to record participants’ gaze during the task at a sampling rate of 300 Hz. For real-time, gaze-contingent control of the task, we used the Tobii Pro SDK to interface with MATLAB. ECG data were recorded using the BIOPAC MP160 system and an ECG100C amplifier (BIOPAC Systems, Inc.) at a sampling rate of 1000 Hz.

### 4.3 Task and Procedure

Upon the arrival, participants were requested to provide informed consent, and disposable electrodes were attached for ECG recordings using a 3-lead configuration. Participants sat 56 cm away from the display with chin rest and performed the “searchlight task,” in which they were required to report the content of each image. Because only the central visual field was made visible, participants had to actively move their eyes to scan the entire image, much like a searchlight. The visible area was limited to a circle with a 40 pixels radius, centered on the foveal region; areas outside this circle were masked in black (Fig. 1). To calculate the coordinates of the visible area, we calculated the average of the coordinates sampled 0.01 milliseconds by the right and left eyes, multiplied each value by the vertical and horizontal dimensions of the display, and then found the median value of the results. We then substituted this value into circle formula to determine the coordinates of the visible area. After 40 seconds of scanning, participants were instructed to verbally report the number of animals and the background context. No feedback was given. The task consisted of 5 practice trials and 21 main trials (one trial per image), which were divided into 3 blocks of 7 trials. A 9-point calibration was conducted before each block, and short breaks were provided between trials. During these short breaks, participants were instructed to remove their chin from the chin rest and sit in a relaxed posture. The total experiment duration was approximately 45 minutes.

### 4.4 Data Analysis

Physiological signal pre-processing was performed using MATLAB 2023a (MathWorks, Inc.). Statistical analysis was conducted using RStudio (RStudio Team, 2023) with R version 4.4.0 (R Core Team, 2023).

#### 4.4.1 ECG Data Analysis

We first applied a 5 Hz high-pass, first-order Butterworth filter on the raw ECG data to remove body-movement-related artifacts. R-wave peaks were then identified as local maxima with a minimum peak distance of 500 msec, and R–R intervals were computed as the differences between consecutive peaks. The analysis was automated with a custom-written MATLAB program and visually inspected for accuracy.

#### 4.4.2 Gaze Data Analysis

Gaze data analysis focused on detecting the timing of events, including saccades, fixations, and blinks. First, we converted the left eye gaze coordinates from pixels to millimeters, then applied an 11-point moving average filter, and performed differentiation to calculate the eye-movement velocity. A saccade event was detected when the velocity exceeded the threshold of 2.5^°^/sec. From the consecutive points meeting this criterion, the point corresponding to the maximal velocity was marked as the saccade time point. Fixations were identified when the velocity was between a 0 and –50^°^ and lasted over 200 msec, with the midpoint of these data points designated as the fixation timepoint. Blinks were detected when consecutive NaN sequences persisted for more than 150 msec, with the onset of these sequences marked as the blink timepoint. Eye movement data that did not exceed these thresholds or criteria were excluded from further analysis.

#### 4.4.3 Circular Analysis

We examined the relationship between oculomotor events and the cardiac cycle by creating a circular representation in which vertices were defined by R–R intervals. The timing of each oculomotor event was calculated as follows:

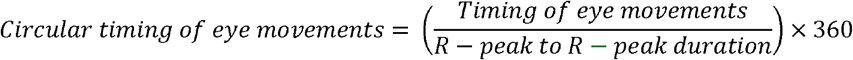

We then obtained a median phase (in degrees) for each type of oculomotor event for each participant. Using each participant’s representative value, we obtained a group-level distribution of the circular timing of the events. To test the uniformity of phase distribution, we employed a Rayleigh test for each oculomotor type separately.

## 5. Acknowledgements

This work was supported by JSPS KAKENHI (Grant #23K24751). We thank T.H. for valuable guidance on data analysis, and K.M. for insightful comments during manuscript preparation.

